# Genetic characterization of ABC pesticide transporters establishes *Drosophila* as an *in vivo* model for toxicogenetics

**DOI:** 10.1101/2022.01.27.477983

**Authors:** Shane Denecke, Hằng Ngọc Bao Lương, Venetia Koidou, Maria Kalogeridi, Rafaella Socratous, Steven Howe, Kathrin Vogelsang, Ralf Nauen, Phil Batterham, Sven Geibel, John Vontas

## Abstract

Pesticides remain one of the most effective ways of controlling agricultural and public health insects, but much is still unknown regarding how these compounds reach their targets. Specifically, the role of ABC transporters in pesticide absorption and excretion is poorly understood, especially compared to the detailed knowledge in mammalian systems. Here, we present a comprehensive characterization of pesticide transporters in the model insect *D. melanogaster*. An RNAi screen was performed, which knocked down individual ABCs in targeted epithelial tissue, examining the subsequent changes in sensitivity to the pesticides spinosad and fipronil. This simultaneously implicated a novel ABC drug transporter *CG4562* but also highlighted a predominant role for the P-glycoprotein orthologue *Mdr65*. Further characterization of the P-glycoprotein family was performed via transgenic overexpression and immunolocalization, finding that *Mdr49* and *Mdr50* play enigmatic roles in pesticide toxicology perhaps determined by their different subcellular localizations within the midgut. Lastly, heterologous expression of the *Mdr65* orthologue from the major malaria vector *Anopheles gambiae* was used to establish an *in vivo* characterization system for the characterization of P-glycoproteins from non-model insects in *D. melanogaster*. This study provides the basis for establishing *Drosophila* as a model for toxicology research regarding drug transporters.

## Introduction

Small molecule pesticides have been the most versatile method of control for both agricultural and public health insect pests. However, there is an urgent need for new compounds due to the emergence of pesticide resistance and unacceptably high deleterious effects on off-target species. One obstacle in the design of new pesticides is bioavailability, or the proportion of a drug that can cross epithelial barriers and reach its target within the body. Although a variety of different factors determine a drug’s ability to cross these epithelia [1], transporters are particularly interesting from a genetic perspective. By translocating molecules across the lipid bilayer, they can either decrease or increase bioavailability depending on their localization and direction of transport. Such drug transporters have been extensively studied in the context of pharmaceutical development; regulatory agencies have specified guidelines for transporter conscious drug design [2], and companies have been founded which focus exclusively on drug transporter services (https://www.solvobiotech.com/). However, the role of these transporters in pesticide toxicology has so far been understudied.

Of particular interest is the ATP-binding cassette (ABC) superfamily. Each functioning ABC transporter is composed of two conserved nucleotide binding domains (NBDs) and two transmembrane domains (TMDs) involved in substrate recognition [3]. Individual transporters are classified into families (A-H) based on homology, which can suggest function but does not allow prediction of which specific substrates are transported by members of a given family. Selected members of the A, B, C, and G families are known to transport xenobiotics in mammalian systems [4]. Perhaps the best studied transporter is P-glycoprotein (aka ABCB1, P-gp), a from the B family. This protein was first identified in a cancer resistant mammalian cell line [5], and has been found to act as a polyspecific drug transporter across a range of different taxa.

Much work on ABCs in insects has been focused on their role in detoxifying pesticides and plant secondary metabolites [6]. Most of this work has been indirect, implicating pesticide transporters via their upregulation in resistant populations or following pesticide exposure. While some transporters have been functionally characterized [7–9], the diversity of transporters that act on pesticides is poorly understood. Like other taxa, P-glycoprotein has been the most studied drug transporter in insects. A family wide expansion of P-gp was previously found in Lepidoptera (moths and butterflies), which may underpin their extreme polyphagy [10]. CRISPR-Cas9 removal and *in vitro* studies have also highlighted their conserved role as drug transporters in this order [11–13]. However, a more detailed characterization of P-gp *in vivo* is lacking.

The study of ABC transporters like P-gp is often difficult *in vitro* despite a wide variety of assays being available [14]. More direct methods of transport such as monolayer cell assays and *in vivo* mice models are more accurate but are relatively expensive and thus not available to all laboratories. The fruit fly *Drosophila* represents a potentially powerful system to study ABC transporters *in vivo* at least with respect to toxins such as pesticides [15]. Apart from its advantages as a genetically tractable model organism, it also has physiologically well characterized epithelial tissues such as the midgut [16], Malpighian tubules [17] and blood-brain-barrier (BBB; [17]). Recent papers have also considered the four P-glycoprotein paralogues in *Drosophila. Mdr65* was localized to the BBB and shown to efflux fluorescent dyes [19]. Deletion of this gene along with the tubule predominant *Mdr49* and midgut predominant *Mdr50* significantly increased sensitivity to a variety of pesticides [20–23]. Strikingly, removal of *Mdr49* and *Mdr50* displayed opposite pesticide responses (more tolerant or more sensitive) depending on which pesticide was used. Similar results were found when RNAi was used to target these two genes [23]. This apparent paradox does not have a proven explanation but may be rooted in the localization of these transporters within tissues and cells. Such differences in localization among the well-studied ABCC (MRP) transporters in mammals have explained such opposing effects [24]. However, these questions have so far not been investigated in insects.

Here we present a series of experiments which sought to use *D. melanogaster* to i) identify novel pesticide transporters ii) characterize their effect on pesticide toxicology *in vivo* and iii) establish *Drosophila* as a heterologous pesticide transporter model.

## Results

### An in vivo RNAi screen identifies putative pesticide transporting ABCs

Of the 56 total ABC transporters in *D. melanogaster*, 37 belong to the B, C, or G subfamilies that have been previously implicated as drug transporters in mammalian systems. For our study we further only considered transporters they had an expression level of >5 RPKM in a given tissue on the FlyAtlas expression database or were present above that threshold in a separate midgut transcriptome [25,26]. This narrowed the list to 33 transporter genes and 59 combinations of transporter tissue combinations tested in this study (Table S1).

An RNAi screen was performed whereby individual transporters knocked down in a single tissue were assessed for their insecticide tolerance. Knock down of several transporters in a single epithelium yielded differences in sensitivity compared to its respective control genotypes, while the majority showed either no difference or were inconsistent (Figure 1; Table S2). Most notably, knock down of *Mdr65* substantially increased the sensitivity of both spinosad and fipronil (Figure 1). To a far lesser extent knockdown (KD) of CG32091, CG3164, and *CG4562* in the midgut tissue increased tolerance to spinosad. Furthermore, KD of CG8799 and *CG9664* in the tubules increased tolerance to spinosad. However, the variation inherent in bioassays and RNAi preclude firm conclusions being drawn from only the RNAi data alone.

**Figure 1:**
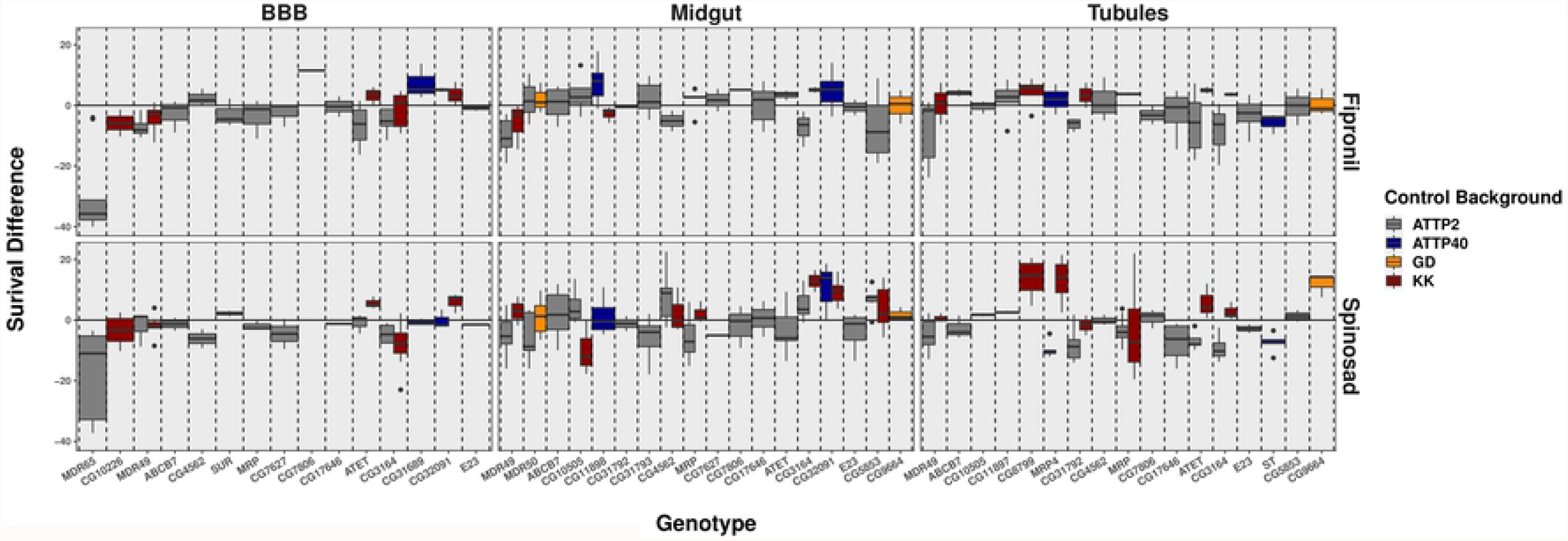
ABC Transporter RNAi screen A knockdown of ABC transporters was performed in the BBB (left), Midgut (middle), and Malpighian tubules (right), followed by a bioassay on either spinosad (bottom) or fipronil (top). Vertical dashed lines distinguish the different ABC transporters targeted by RNAi. Horizontal solid lines signify how the genetically matched control line behaved, while boxes reflect the degree of difference between the knockdown and control. In other words, the further from the vertical line a box is the more susceptible (lower) or resistant (higher) the knockdown was. Boxes are color coded based on their genetic background. From Bloomington TRIP stocks, AttP2 was grey, AttP40 was blue. From the VDRC, GD was orange and KK was red.

### CRISPR-Cas9 of uncharacterized ABC transporter increases spinosad tolerance

*CG4562, CG3164*, and *CG32091* were selected for knockout (KO) using CRISPR-Cas9 as they showed the most consistent differences in the RNAi toxicology screen. A KO line for *CG3164* was unable to be established as it did not homozygose from the balancer stock, suggesting the gene plays an essential role. KO of *CG32091* and *CG4562* were successful. Of these, only *CG4562* showed a decrease in spinosad sensitivity which mimicked the RNAi phenotype (Figure 2; Table S3). This suggests that *CG4562* plays a role in spinosad transport in *D. melanogaster*, but the magnitude of this change was far less than that observed for *Mdr65* knockdown.

**Figure 2:**
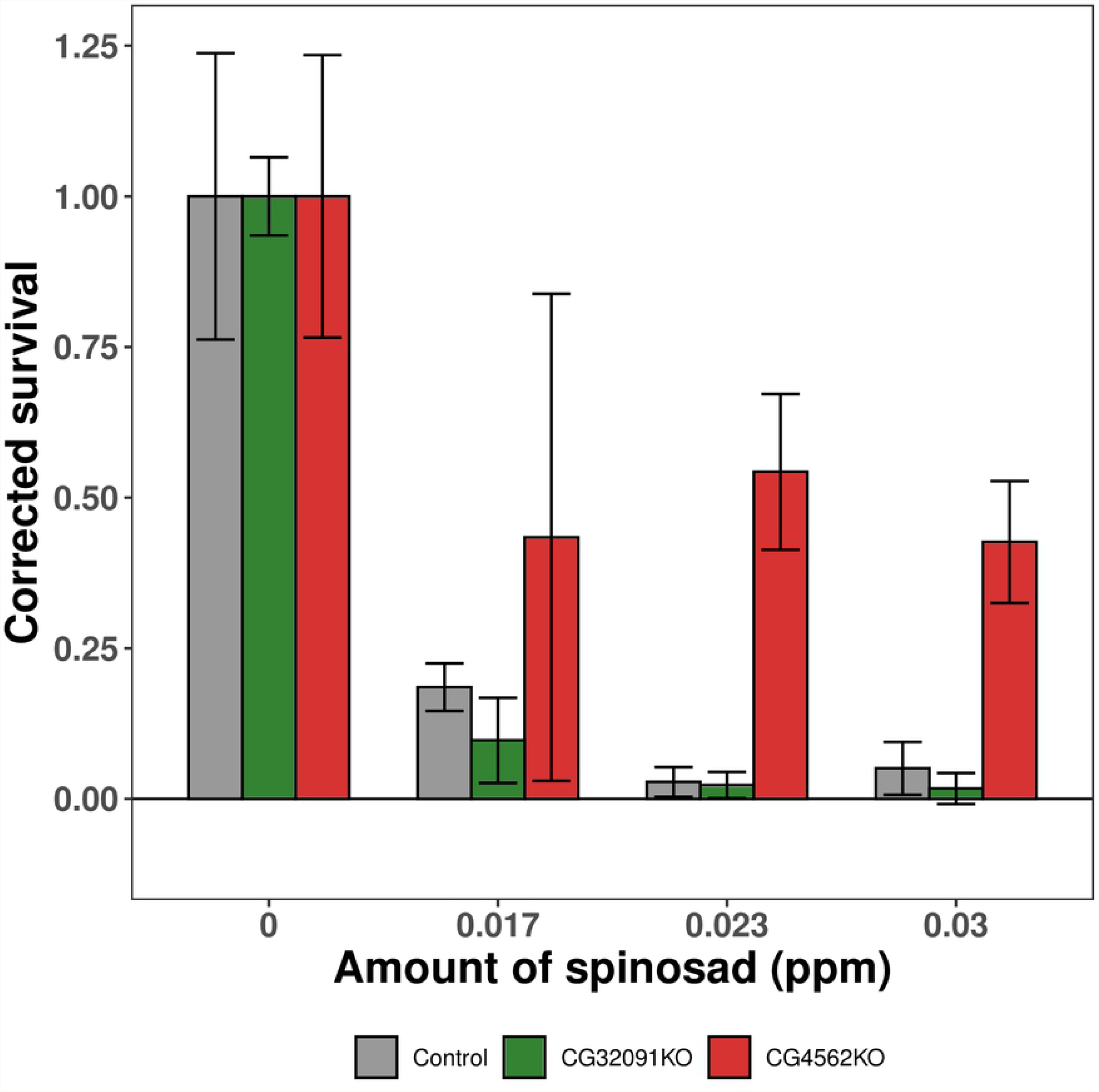
CRISPR-KO of candidate drug transporters Bioassay data from the KO of CG32091 and *CG4562* is shown. The genetic control background (Nanos-Cas9) and the CG32091 showed a strong response, indicated by a low survival proportion on the y-axis. Removal of *CG4562* showed a higher proportion of survival across all doses of spinosad (x-axis). Error bars represent 95% confidence intervals generated by Schneider-Orelli’scorrection.

### Overexpression of P-glycoprotein in Drosophila

In the RNAi screen *Mdr65* knockdown provided the most consistent impact on the response to spinosad and fipronil across the three tissues tested (Figure 1). Further, CRISPR-Cas9 based KO of *Mdr65* and its paralogues *Mdr49* and *Mdr50* indicates that each of the genes has a role in insecticide response [20]. We thus sought to expand upon these findings using transgenic UAS responder lines containing the ORFs of *Mdr49, Mdr50* and Mdr65. Overexpression of *Mdr65* at the BBB increased tolerance to spinosad and showed no difference against fipronil in contrast to the striking increase in susceptibility observed with RNAi in this tissue (Figure 3A,B). Very minor differences in response were observed when *Mdr49* or *Mdr50* were driven in their respective tissues of the tubules or the midgut respectively (Figure 3 C-F; Table S4).

**Figure 3:**
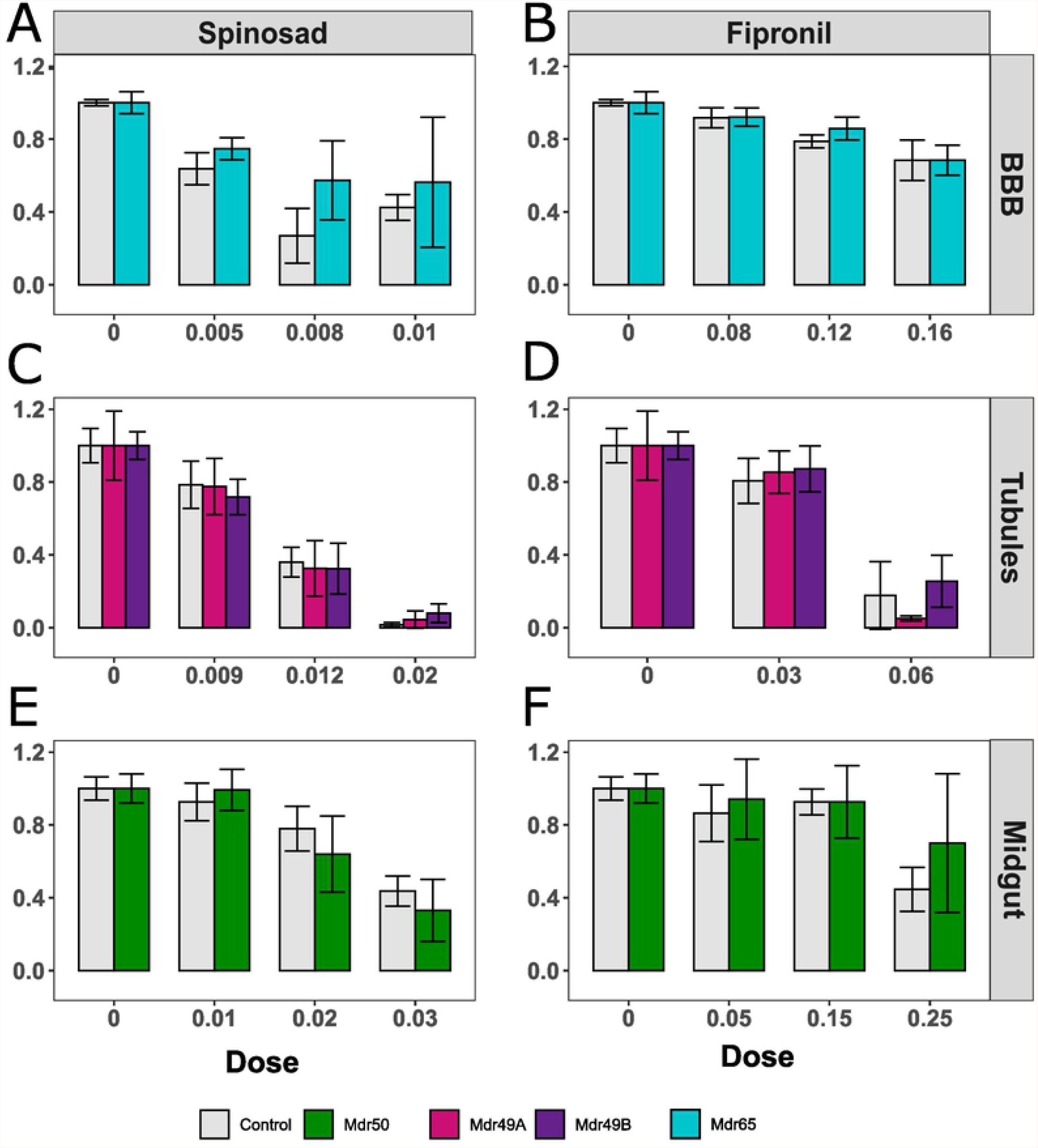
Tissue specific overexpression of P-gp paralogues Bioassay data is shown for both spinosad and fipronil against genotypes selectively overexpressing a single P-glycoprotein orthologue (*Mdr49*; C-D, Mdr50; E-F, or Mdr65; A-B) in the specific tissue corresponding to their proposed localization. Grey bars indicate genetically matched controls in all samples while colors correspond to specified overexpression genotypes.

Transgenic expression of these P-gp genes with a strong ubiquitous driver yielded a larger effect. Both *Mdr49A* and *Mdr65* increased sensitivity to spinosad while *Mdr50* overexpression lines were slightly more resistant (Figure 4B,D; Table S5). With the pesticide nitenpyram ubiquitous expression of both *Mdr65* and *Mdr50* caused resistance while overexpression of *Mdr49* caused increased sensitivity (Figure 4 A,C). Overexpression of *Mdr49B* had limited effect. These results agree with previous findings [20,23], demonstrating that global deletion or expression of genes can have variable effects on the direction of toxicological change (more susceptible or resistant) depending on the gene and pesticide under investigation.

**Figure 4:**
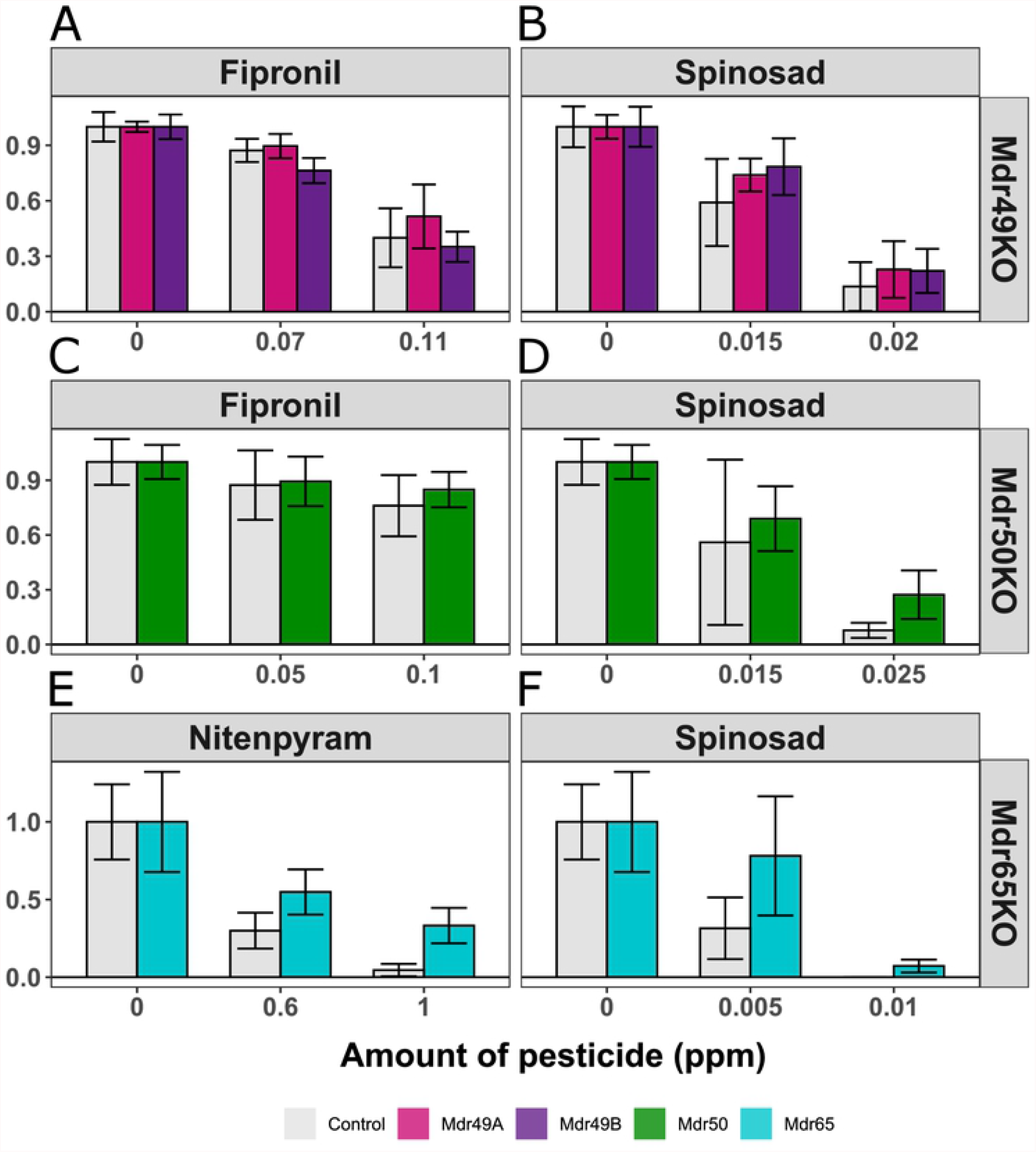
Global overexpression of P-gp paralogues Bioassay data is shown for the pesticides spinosad (B,D) and nitenpyram (A,C), the latter chosen due to the difficulty of scoring fipronil toxicology in this genetic background. As the *Mdr65* overexpression genotype was generated in a distinct background it was considered separately (A-B), while all other overexpression constructs were analyzed jointly (C-D). In each panel the y-axis represents survival while the x-axis signifies the dose of the pesticide under investigation.

Rescue experiments whereby we replaced an endogenous copy of P-gp with itself in a specific tissue. The expression of *Mdr49A* or *Mdr49B* in the Malpighian tubules of the Mdr49KO background did not have any measurable effect on the toxicology of spinosad or fipronil (Figure 5 A,B; Table S6). Likewise, the reintroduction of *Mdr50* into the midgut increased survival but only very subtly; the effect was observed only at one dose (Figure 5 C,D). A much clearer effect was seen with the rescue of *Mdr65*. Significantly higher survival was observed in both pesticides when a functional copy of *Mdr65* was added into the Mdr65KO background specifically at the BBB (Figure 5 E,F). These results highlight the subtlety of the effect seen with *Mdr49* and *Mdr50*, but also suggest the *Mdr65* rescue system as a viable way of detecting transgene function *in vivo*.

**Figure 5:** Genetic rescue of *Drosophila* Knockouts Genetic rescue of the three knockouts Mdr49KO, Mdr50KO, and Mdr65KO was tested for their pesticide toxicological response. Spinosad (B,D,F), Fipronil (A,C), and Nitenpyram (E) were used against different genotypes, with the latter being chosen due to the difficulty of scoring fipronil toxicology in this genetic background. In each panel the x-axis represents pesticide dose and the y-axis represents survival corrected with abbots correction. Error bars represent 95% confidence intervals generated by Schneider-Orelli’s correction.

### Immunolocalization of endogenous P-gp

Given that overexpression of P-glycoprotein orthologues in *Drosophila* yielded qualitatively different phenotypes depending on the insecticide and gene in question, we sought to localize these proteins with immunofluorescence. The C219 antibody was used, which targets an epitope common to all P-glycoprotein paralogues found in *Drosophila*, and a genotype with a wild type P-gp gene (Myo-GAL4) was used in comparison with all three P-gp KOs. No staining was found in the fat body in any genotype under investigation, suggesting that this tissue does not contain notable P-gp expression (Figure S1). The dissected central nervous system showed strong staining in all genotypes except Mdr65KO, confirming that the MDR65 protein is the predominant P-gp orthologue at the BBB (Figure S2). Removal of *Mdr49* had a drastic effect and abolished almost all tubule expression especially that concentrated in the lumen (Figure S3 C’). The remaining non-luminal P-gp is presumably due to expression of *Mdr50* or *Mdr65* present in small quantities in the tubule.

Lastly, the midgut was examined which provided the most complicated yet interesting tissue under study. In wild type and Mdr65KO flies, P-glycoprotein was found localized primarily to a region of the posterior midgut (defined as R5 by [27]), which matches transcriptomic data for these genes (Figure 6 A-B). There were also lower amounts of expression detectable in the anterior midgut near the gastric caeca. However, Mdr50KO and Mdr49KO genotypes appeared to show opposing subcellular localizations. While Mdr50KO showed an apical specific phenotype, Mdr49KO localized only to the basolateral side (Figure 6 C-D). This suggests that different paralogues differ by their subcellular localization despite residing in the same tissue.

**Figure 6:** Immunolocalization of P-glycoprotein paralogues Immunostaining is shown for the control genotype (Myo; A) and Mdr65KO (B), Mdr49KO (C), and Mdr50KO (D). For each row, zoomed out images of the whole midgut are shown along with zoomed images based depending on where fluorescence was detected. Brightfields are shown along with confocal images and merges for each column. In each image, DAPI stained nuclei are in blue, phalloidin stained actin is in green, and the C219 antibody localization (P-gp) is in red.

### Transgenic Expression of P-glycoprotein from a malaria vector

We lastly sought to extend our findings to P-glycoproteins from pest species such as the malarial vector *Anopheles gambiae*. Strong, ubiquitous expression of *AgP-gp* (AGAP005639) in *Drosophila* resulted in increases in sensitivity to both nitenpyram and spinosad with effect sizes on par with or greater than the endogenous *Drosophila* proteins (Figure 7; Table S5). Given the larger effect sizes observed when rescuing the Mdr65KO at the blood brain barrier, *AgP-gp* was also expressed specifically at the blood brain barrier in an Mdr65KO background this suggested a higher tolerance to the pesticide spinosad, but not fipronil (Figure 8; Table S7). The expression of *Mdr49A* or *Mdr49B* did not yield any effect in this system suggesting that different paralogues may not be active in this tissue.

**Figure 7:** Global overexpression of P-glycoprotein in *A. gambaie* The single P-glycoprotein orthologue from Anopheles gambiae was transgenically expressed in *D. melanogaster* under the control of a strong ubiquitous promoter (Actin-GAL4). These transgenic flies (dark grey) were then compared with their matched controls (light grey) using a larval toxicology assay against Nitenpyram (left) or Spinosad (right). In each panel the x-axis represents pesticide dose and the y-axis represents survival corrected with abbots correction. Error bars represent 95% confidence intervals generated by Schneider-Orelli’s correction.

**Figure 8:** Genetic rescue of *Drosophila* Mdr65KO with *AgP-gp* Three different UAS-P-gp constructs expressing *Mdr49A* (Pink), *Mdr49B* (Purple) and *AgP-gp* (dark grey) were expressed in a Mdr65KO background and tested on the pesticides nitenpyram (left) and spinosad (right). In each panel the x-axis represents pesticide dose and the y-axis represents survival corrected with abbots correction. Error bars represent 95% confidence intervals generated by Schneider-Orelli’s correction. Although there is no established statistical test for Schneider-Orelli’s correction non-overlapping 95% confidence intervals are considered a very conservative estimate and significance stars were included in this graph reflecting this.

## Discussion

The emergence of resistance and unacceptably high externalities posed by the current generation of pesticides necessitates novel, safe, and effective compounds for insect control. Bioavailability presents an obstacle to rational pesticide design, and this is worsened by a poor understanding of insect drug transporters *in vivo*. Here, we develop the *Drosophila* model system for transporter mediated toxicology by i) characterizing the diversity of pesticide transporters in *Drosophila* ii) providing a more in-depth characterization of P-glycoprotein and iii) heterologously expressing an orthologous mosquito transporter in *D. melanogaster*.

### The diversity of ABCs involved in pesticide transport

One of the goals of this study was to identify which ABCs were involved in pesticide transport, and a toxicology screen of RNAi lines was performed to address this. Identification of the previously known *Mdr65* gene suggested that the assay could detect true positives and the screen further suggested a previously uncharacterized gene (*CG4562*) which transports spinosad in the midgut (Figure 1). This was confirmed by CRISPR-Cas9 KO, which phenocopied the RNAi knockdown (Figure 2). *CG4562* belongs to the ABCC family, several members of which have been implicated in drug transport in humans [28] and other insects such as the human body louse [29], red flour beetle [8] and even in *D. melanogaster* [30]. Unlike several other ABCC drug transporters identified in insects, removal of *CG4562* increases tolerance to spinosad rather than increasing sensitivity. This runs counter to the narrative of ABCs as “resistance” genes and highlights a more complex role for transporters in influencing the distribution of compounds around the body. Similar findings in mammalian ABCC genes have shown that transporters localizing to the basolateral side of tissues like the intestine have the effect of pumping compound into the body [24]. While *CG4562* was not localized in this study, we hypothesize it to be present at the basolateral membrane and thus pump toxin out of the midgut cell into the hemolymph.

However, several limitations of the assay may have skewed a bias of the RNAi screen towards false negatives. For example, *Mdr49* and *Mdr50* were not detected in the RNAi screen despite CRISPR-Cas9 KO yielding clear effects [20]. This may stem from considering each tissue in isolation or may also represent a limitation of the strength of the knockdown achieved in this study, either of which could mask subtle effect sizes. Future studies could make use of the next generation of genetic tools in *Drosophila* such as somatic CRISPR for tissue specific removal or simply making heritable mutations in a larger range of transporters [31] which would likely elicit larger effects. Combinatorial knockouts or knockdowns may also be an option as synergistic action has been previously associated with toxicology in both mammals [32] and insects [33].

### P-glycoprotein plays a predominant role in insecticide toxicology

Knockdown of the previously known *Mdr65* gene at the blood brain barrier created an effect size that was an order of magnitude higher than for any other transporter, suggesting a predominant role for this protein in pesticide toxicology. Such a role would match the situation in mammals where P-glycoprotein has by far the largest effect in most tissues [28]. Interestingly, P-glycoprotein has only been rarely associated with insecticide *resistance*, defined as the abnormally high levels of tolerance to pesticides causing control failures. Selected studies have shown P-gp’s role in resistance [34], but this mechanism does not appear to be widespread among insects compared to other methods like target site mutations or upregulation of drug metabolizing enzymes [35]. Indeed, in the current study only strong ubiquitous expression was able to generate changes in toxicology and these were modest compared to the effect of removing the gene (Figure 1, Figure 3). P-glycoprotein thus appears to play a strong role in the toxicology of pesticides, but increased expression may have only a limited effect on pesticide resistance. Interestingly, an opposite role is played by some cytochrome P450s; overexpression of *Cyp6g1* confers insecticide resistance, while removal has a limited effect [36].

Strikingly, the ubiquitous expression of *Mdr49* or *Mdr50* caused opposite changes in toxicology (increased tolerance or increased sensitivity) depending on both the pesticide and transgene, agreeing with previous CRISPR-Cas9 data [20,23]. These differences may be due to changes in localization, and immunostaining at the cellular level indicated a change in subcellular localization of *Mdr49* and *Mdr50* genotypes (Figure 6). However, it is currently unclear how any potential localization changes cause the observed changes in toxicology among pesticides. Another hypothesis is that a compensatory mechanism exists whereby the removal of one transporter impacts another as was seen for organic anion transport in the tubule [37]. Lastly, there may be restrictions as to which tissue a given paralogue can act. Ubiquitous expression of different paralogues yielded different effects (Figure 4), and only certain paralogues *(Mdr65* and *AgP-gp*) were shown to be functional in the BBB rescue system (Figure 8). Future work will be needed to address this paradox, which promises to have important implications for toxicokinetics and for basic research on how transporter gene’s function.

### Drosophila as a model system for ABC mediated toxicology

The characterization of the P-gp orthologue from *A. gambiae* showed that it was able to influence the toxicological profile of the pesticide spinosad when expressed both ubiquitously and in a tissue specific *Mdr65* rescue system at the BBB (Figure 7,8). Although this system no doubt requires optimization, the successful heterologous expression of a pest transporter *in vivo* opens the door for the study and comparison of these genes in the future. So far it has been challenging to find suitable expression systems for drug transporters and *Drosophila* represents an unexplored alternative. Side by side expression of transporters from pest species (e.g., mosquitoes, agricultural pests) with non-target organisms (e.g., humans, bees), in a *Drosophila* like system could provide the basis for pest selective insecticides in the future. Such heterologous expression in *Drosophila* has been used to previously characterize pesticide target proteins [38], and given the challenging interpretation of *in vitro* transporter data [14], *Drosophila* may provide a welcome alternative.

## Materials and Methods

### Previously generated Drosophila genotypes

Several driver genotypes were used in this study which expressed GAL4 in defined epithelial tissues (Luong et. al under review). The Mex-Gal4 line was previously reported (Phillips and Thomas, 2006) and drove expression specifically in the midgut. Similarly, the Myo1A GAL4 driver (Myo-GAL4; 67057) was used as an alternative to Mex-G4 in the P-gp rescue experiments as it also drives expression in the midgut enterocytes. The Urate_Oxidase-GAL4 (UO-GAL4) localized expression specifically in the Malpighian tubules [39]. Lastly, a previously published driver which localized GAL4 to the subperineural glial cells of the BBB was used (SPG-GAL4; [40]. Strong ubiquitous expression was also achieved with Actin-GAL4 (Bloomington #3953).

Publicly available UAS-RNAi lines were obtained from the transgenic RNAi project (Trip) or the Vienna *Drosophila* RNAi Centre (VDRC) and contained RNA interference sequences targeting an individual ABC transporter under the control of a UAS promoter (Table S1; [41,42]. Crossing of any one of these flies to a GAL4 would express the RNAi construct and knock down the specified ABC in the specified tissue.

### Generation of CRISPR-Cas9 knockout genotypes

Selected genes identified in the RNAi screen were confirmed by removing the majority of the gene from the genome using CRISPR-Cas9. Pairs of sgRNAs for each were first designed via the CRISPR optimal target finder [43]. These were selected so that one sgRNA targeted a sequence at the 5’ and 3’ end of the gene. Each sgRNA was cloned into the Bsbi site in the pU6 vector by annealing overlapping custom primers (Table S2). Pairs of pU6-sgRNA plasmids were then injected into fly lines expressing Cas9 under the control of the Nanos promoter either on the 2nd or 3rd chromosome (Bloomington # 78781 & 78782). In either case the chromosome carrying Cas9 was crossed out so that all deletions were eventually in identical backgrounds apart from the ABC transporter being removed.

### Transgenic expression genotypes

Transgenic lines containing an open reading frame (ORF) of a given transporter regulated by a GAL4 Inducible UAS promoter were created. ORFs were first amplified with gene specific primers and cloned into the pGEM-T-easy shuttle vector. The fragments were then digested with flanking NotI sites, subcloned into the NotI site of the pUASt-AttB vector [44], and injected into the AttP40 (Bloomington #25709) or the VK13 (Bloomington #24864) *D. melanogaster* lines harboring AttP sites and an endogenous source of φC31 integrase. This strategy was used to introduce *Mdr49A, Mdr49B, Mdr50, Mdr65*, and the P-gp derived from the VK7 stain of Anopheles gambiae (*AgP-gp*).

In addition to expression in “wild type” backgrounds, transgenic expression of these P-gp genes was performed in P-gp KO backgrounds [20] in order to observe any complementation (rescue) of the initial toxicology phenotype. In order to match known endogenous expression patterns, *Mdr49* was expressed specifically in the Malpighian tubules, *Mdr50* specifically in the midgut, and *Mdr65* specifically in the BBB. For *Mdr49* and *Mdr50*, stable lines were made carrying i) the deletion ii) the UAS-transgene and iii) the GAL4. As the KO and transgenic GAL4 cassette and KO was on the same chromosome recombination was used to generate flies carrying both alleles. Each of the two previously characterized *Mdr49* isoforms (A and B; [21] were considered separately. For *Mdr65*, the transgene and the BBB-GAL4 cassette (on the 2nd chromosome) were both independently crossed into the Mdr65KO background separately and finally crossed together to make the rescue line. For the heterologous BBB rescue system, the UAS-Pgp construct was inserted into the VK13 landing site (3rd chromosome) and was recombined with the Mdr65KO allele (also 3rd chromosome) and crossed into a background with BBB-GAL4 on the 2nd. Due to genetic incompatibilities irrespective of the genes under investigation, BBB-GAL4 was unable to be homozygosed with the Mdr65KO 3rd chromosome, so all rescue lines were maintained and assayed over a 2nd chromosome CyO balancer.

### Insecticide bioassays

Fipronil (CAS# 120068-37-3) and Spinosad (CAS# 168316-95-8) were both obtained as analytical standards from Sigma. Insecticide bioassays were performed as described previously [20]. First instar larvae were obtained by collecting between 150-200 females and 100 males of defined genotypes and allowing them to lay eggs on cherry juice agar plates for 24 hours. After this period, the plates were changed, and the eggs washed onto a fine mesh. Eggs were then transferred onto fresh cherry juice agar plates and left undisturbed for an additional 24 hours which generated a population of 1^st^ instar larvae. 50 larvae were transferred to standard media dosed with a defined concentration of insecticide. Survival was scored after 16 days by counting the number of total pupae in the vial. For genotypes carrying a CyO balancer, 100 larvae were picked/vial and total numbers of straight wing flies were counted as there were no significant differences among the number of CyO emerging individuals (data not shown). Raw mortality data was then analyzed in the R statistical environment.

### Immunolocalization of P-glycoprotein

In order to analyze protein localization of C219 P-gp, immunostaining was performed as described previously [19], with some modifications. In brief, third instar larvae were dissected in SF-900 II insect medium (SFM, Cat. No 10902-096, Gibco) enriched with penicillin-streptomycin mix (50x, Cat. No P4458, Sigma-Aldrich) and placed on ice. Following dissections, the tissues were incubated in 4% formaldehyde solution (Methanol free, Thermo Scientific, 28906) for 60 min and then in 3% Blocking Solution (BS, 3% BSA, 0.1% Triton in PBS1X) overnight at 4°C. Tissues were incubated with mouse anti-P-glycoprotein C219 antibody at 3.6μg/mL (Glycoprotein Monoclonal Antibody C219, Cat. No MA1-26528, Invitrogen) overnight at 4°C. On the third day the tissues were incubated with Alexa Fluor 555-conjugated anti-mouse IgG secondary antibody (Invitrogen) (1:1,000 in BS) for 1.5 hours at RT. After BS washes, actin filaments were stained with Alexa Fluor 488-conjugated phalloidin (1:50 in BS for 30 min at RT, Invitrogen Thermo Scientific, A12379) and nuclei were stained with DAPI (1:100 in BS for 20 min at RT, AppliChem, Cat. No A1001). Tissues were then gently rinsed twice with PBS1X, prior to mounting in Vectashield (Vector Laboratories, H-1000-10) on SuperFrost+ slides. Imaging was conducted using a Leica TCS SP8 confocal laser scanning microscope unit, equipped with an inverted microscope (DMI6000 CS), a high-end Scientific fluorescence CCD camera (Leica DFC365FX) and a set of the appropriate high quality lasers, housed in the Institute of Molecular Biology and Biotechnology (IMBB) Microscope Facility of Foundation for Research & Technology-Hellas (FORTH).

## Acknowledgments

The authors would like to thank Ioannis Livdaras for performing *Drosophila* microinjections. This work was funded as a component of a joint collaboration between Bayer CropScience and the Institute of Molecular Biology and Biotechnology (Greece). Shane Denecke and Ngoc Bao Hang Luong were funded with this collaboration. Kathrin Vogelsang, Ralf Nauen, and Sven Geibel are employees of Bayer CropScience.

## Figure/Table Legends

**Figure S1:**
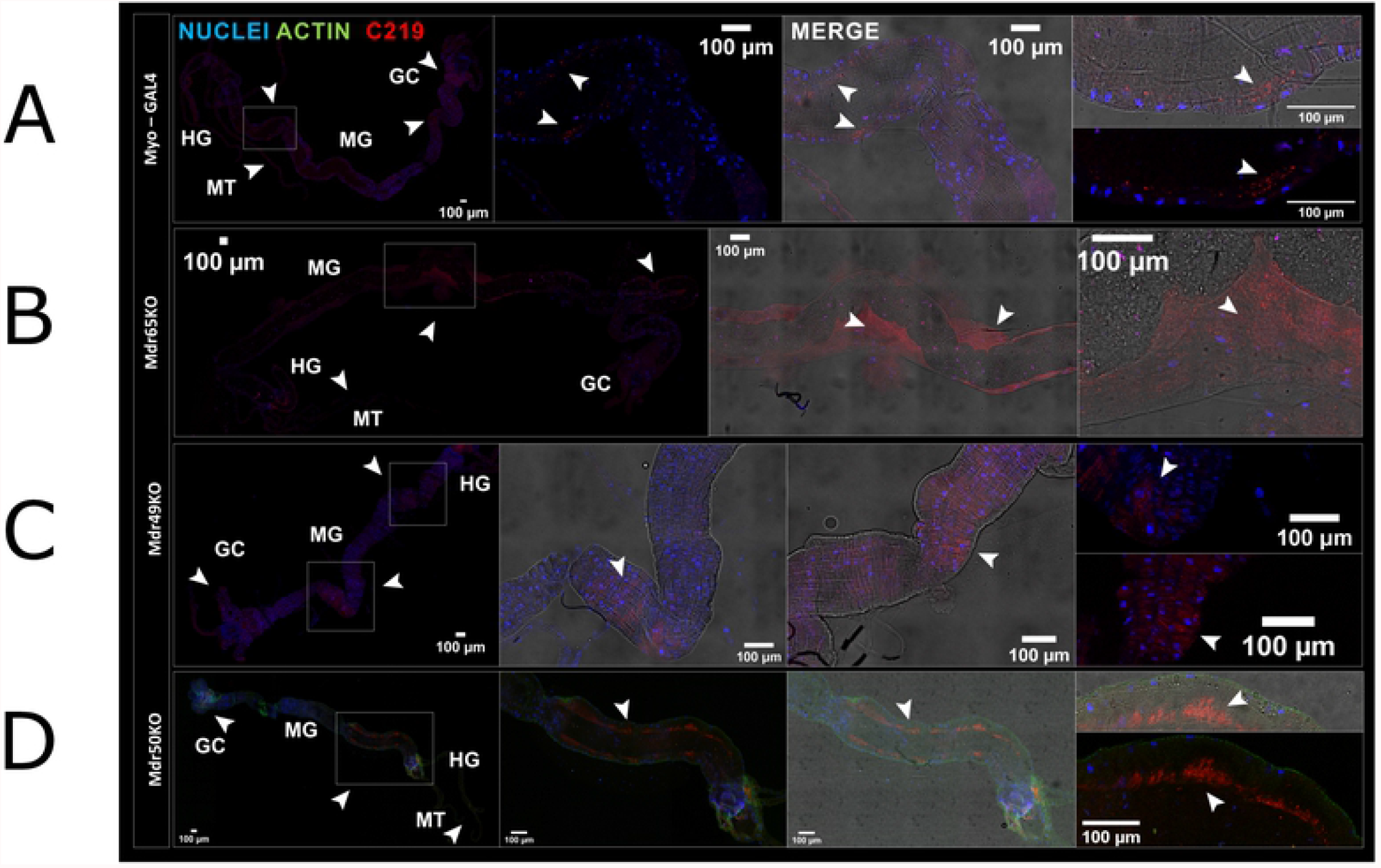
P-gp staining of the *Drosophila* brain The central nervous system was stained was stained for the wild-type control (Myo-GAL4; Top row) along with Mdr65KO (2^nd^ row down), Mdr49KO (3^rd^ row down), Mdr50KO (bottom). Blue indicates nuclei while red indicates the C219 P-gp signal.

**Figure S2:**
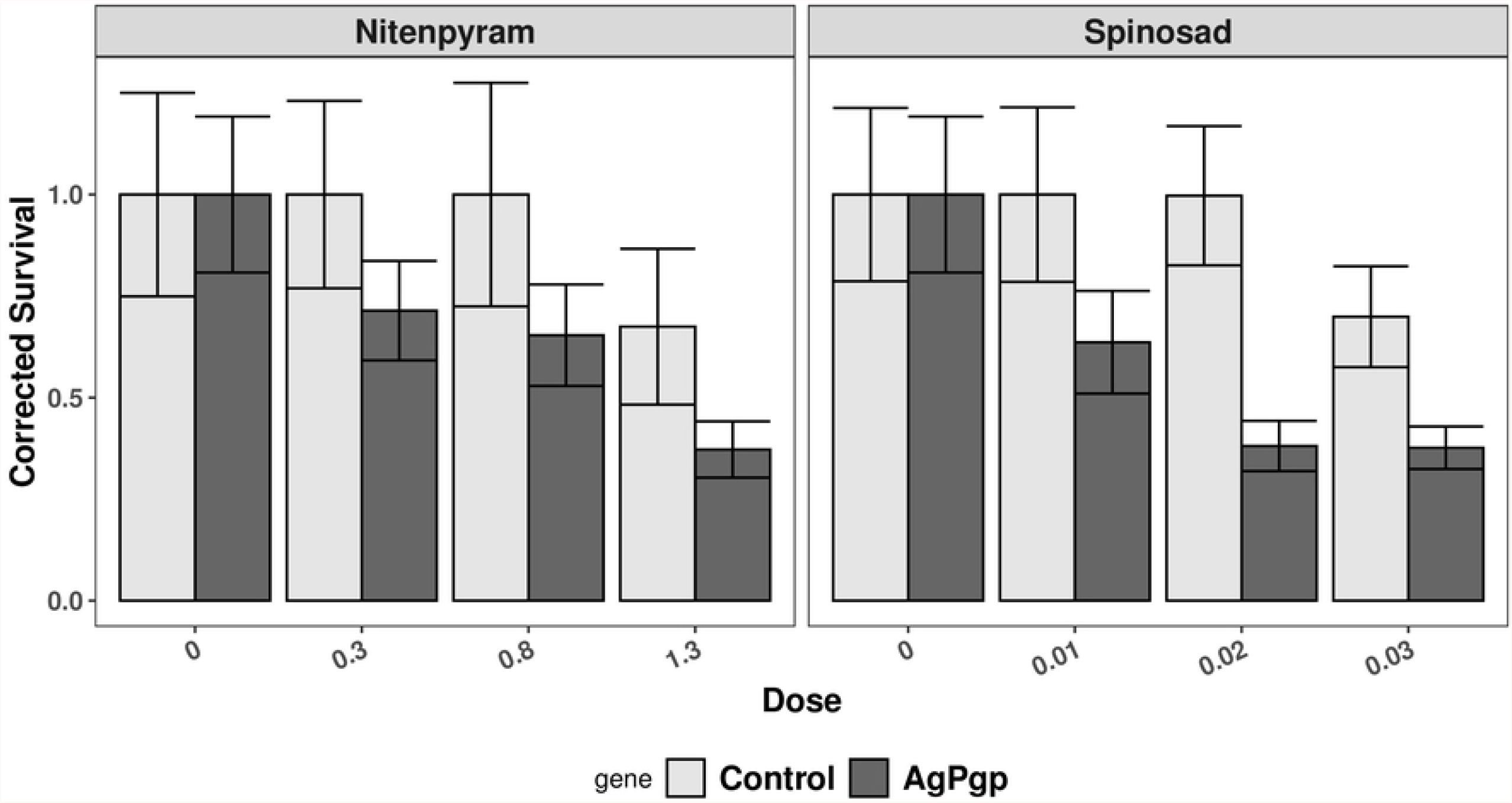
P-gp staining of the *Drosophila* fat body The fat body was stained was stained for the wild-type control (Myo-GAL4; Top row) along with Mdr65KO (2^nd^ row down), Mdr49KO (3^rd^ row down), Mdr50KO (bottom). Blue indicates nuclei while red indicates the C219 P-gp signal.

**Figure S3:**
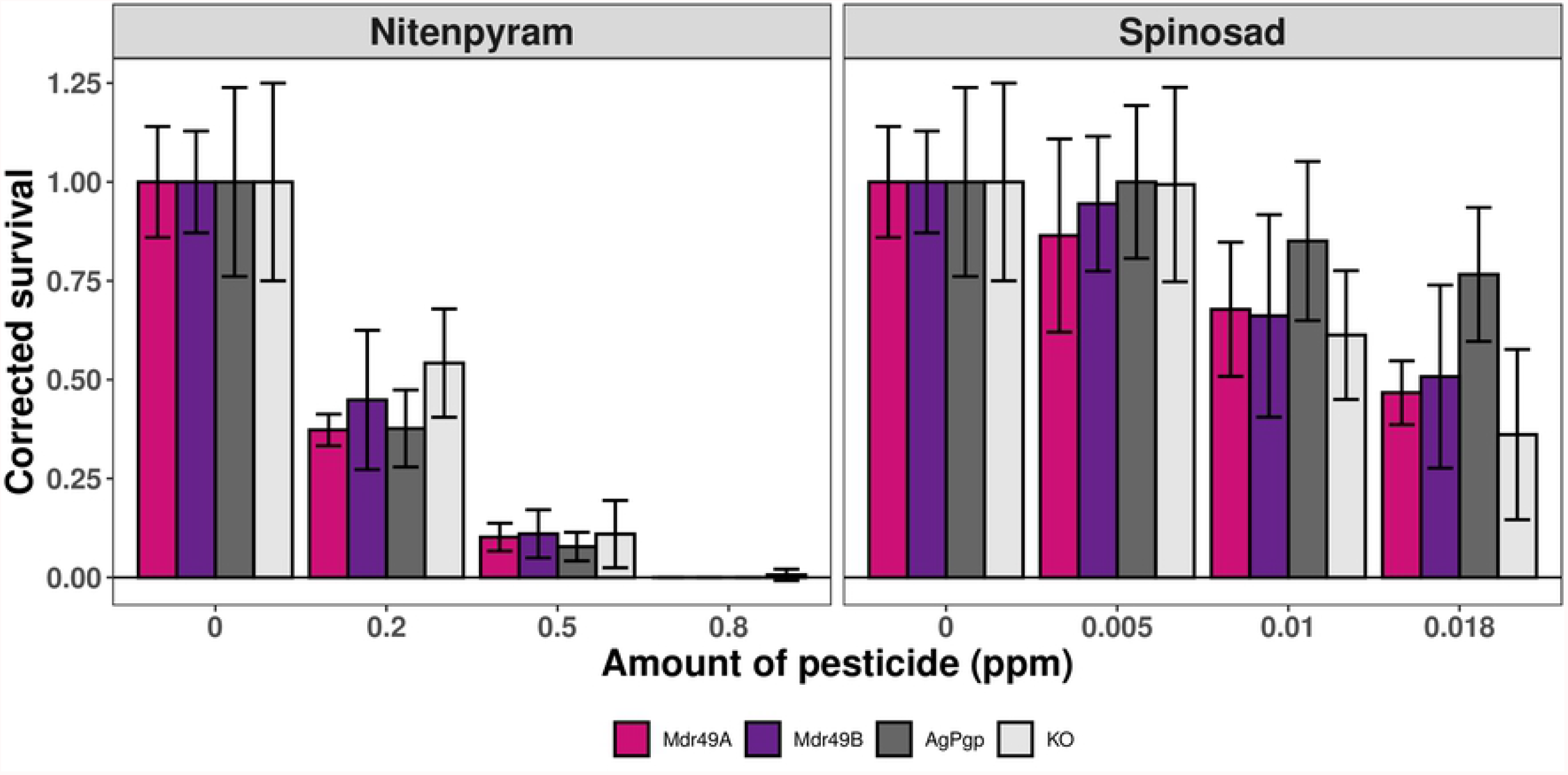
P-gp staining of the *Drosophila* Malpighian tubule The Malpighian tubule was stained was stained for the wild-type control (Myo-GAL4; Top row) along with Mdr65KO (2^nd^ row down), Mdr49KO (3^rd^ row down), Mdr50KO (bottom). Blue indicates nuclei while red indicates the C219 P-gp signal. Green indicates actin.

